## Supplemental File 1 for "Local density determines nuclear movements during syncytial blastoderm formation in a cricket"

*Supplemental Information*

Seth Donoughe[1,2,*], Jordan Hoffmann[3], Taro Nakamura[1,4], Chris H. Rycroft[3,5,*], Cassandra G. Extavour[1,6,*]

[1] Department of Organismic and Evolutionary Biology, Harvard University, Cambridge, MA, USA
[2] *Present address:* Department of Molecular Genetics and Cell Biology, University of Chicago, Chicago, IL, USA
[3] John A. Paulson School of Engineering and Applied Sciences, Harvard University, Cambridge, MA, USA
[4] *Present address:* Division of Evolutionary Development, National Institute for Basic Biology, Okazaki, Japan
[5] Computational Research Division, Lawrence Berkeley Laboratory, Berkeley, CA, USA
[6] Department of Molecular and Cellular Biology, Harvard University, Cambridge, MA, USA

6 Code, data, and materials availability 13

References 13

### 1 Processing, segmenting, and tracking microscopy datasets

The *Methods* section of the main text includes information on animal culture, collecting embryos, mounting samples, and microscope settings. This section contains details on how we processed datasets for analysis.

#### 1.1 Datasets from lightsheet microscopy

We fused multiview lightsheet datasets with the Multiview Reconstruction plugin[1,2] for Fiji[3,4] (version 2.0.0-rc-30 through 2.1.0/1.5.3). Fluorescent beads (ThermoFisher F8821) embedded in the agarose served as fiducial markers for registration; we used them to align and combine multiple views as a weighted average fusion[1]. We deconvolved datasets for ten iterations with the Multiview Reconstruction plugin[2] on a 32-core workstation with 128 GB of RAM running Ubuntu 14.04.

We performed nucleus segmentation and automated tracking with Ilastik[5] (versions 1.1.5 to 1.3.0). We imported all data as a 4D sequence, manually trained a pixel classifier to differentiate nuclei from background, and identified nuclei using the object classification tool. Then we used Ilastik's automated tracking tool to track all nucleus movements. We used a custom Mathematica script to convert the automated tracking output from Ilastik into an XML file that was parsable by MaMuT[6], the semi-automated tracking plug-in for Fiji[3]. The script to generate the XML file, as well as those needed run the simulations in this study, are available on the GitHub repository that accompanies this paper (`https://github.com/hoffmannjordan/gryllus_nuclear_movements`). Note: the ability to directly transfer object tracks from Ilastik to MaMuT is now included in Ilastik (version 1.3.3). Last, we used MaMuT to manually identify each division and stitch together the Ilastik-generated tracks, which resulted in continuously tracked lineages. For embryos mounted in this manner, approximately two-thirds of the radial depth of the embryo could be imaged clearly (as measured from the surface to a center line connecting the anterior and posterior poles of the embryo), while the signal from the inner-most portion of the embryo was diffused by the yolk of the embryo.

#### 1.2 Datasets from epifluorescence microscopy

We recorded multiple embryos at a time by tiling across a field of microwells, each of which held a single embryo, oriented laterally, following previously described methods[7]. We used Ilastik to automatically track nucleus movements and manually identified division in MaMuT. For embryos mounted in this manner, approximately one-half to one-third of the $z$-depth of the embryo could be imaged clearly, while the signal from the rest of the embryo was diffused by the yolk of the embryo. Thus, the number of well-segmented nuclei fluctuated slightly between time points, depending on the particular paths traveled by individual nuclei; the effect of this variability was strongest for the first few division cycles. Therefore, for two-dimensional timelapse (2D+T) datasets, we performed automated analysis on these datasets beginning at the earliest time point when there were at least 25 nuclei.

#### 1.3 Datasets from confocal microscopy

We included three-dimensional timelapse (3D+T) datasets of embryos laid by females with Act-mtdT and Act-H2B-EGFP transgenes[8]. We used a magnification such that approximately one-fifth of the length and one-half of the breadth of an embryo was recorded at a time. We cropped and assembled these into figures using Fiji. Details of the genotypes of these transgenic lines are included in the main text Methods section *Transgenics and animal culture*.

#### 1.4 Uses of each type of microscopy data

*Multiview lightsheet datasets of blastoderm formation:* These 3D+T datasets enabled us to reconstruct nucleus movements and divisions in detail. We recorded these datasets one at a time. After a syncytial blastoderm had formed, we removed the embryo from its agarose column and cultured it in an incubator as described in the main text Methods section *Collecting and culturing embryos*. Once the embryo hatched, we continued with processing and tracking the nuclei. We tracked and processed four such 3D+T datasets. Figures 1B, 1D, 2A-E, and 3A-H in the main text show data from a single such embryo (the same embryo in all cases). We followed the same procedures and repeated the same analyses on three additional lightsheet datasets. In all cases we found the same patterns shown in Figures 1, 2, and 3: a positive association between cycle duration and nuclear density, biphasic speeds within each cell cycle, a negative association between speed and density in Phase A, and a tendency for Phase A nuclei to move into nearby open space when they are in the interior of the embryo (data not shown).

*Epifluorescence datasets of blastoderm formation:* We also recorded 60 additional 2D+T datasets. Three of these datasets are included in Fig. 5 in the main text as unmanipulated controls to compare to the physically constricted embryos.

*Epifluorescence datasets of constricted embryos:* We recorded 2D+T datasets of physically constricted embryos, imaged using epifluorescence. These data are shown in Fig. 5 of the main text. In our hands, constricted embryos did not hatch, and instead they arrested partway through development (in some cases before a blastoderm had formed, and in others afterwards). The furthest extent of development we observed was just before katatrepsis occurred[9]. Our criteria for including a constriction dataset in the data analysis were as follows: no visible ruptures of the eggshell, no cessation of nucleus movements before the syncytial blastoderm was formed, and no nuclear aggregations during the developmental period under study. In our recordings of unmanipulated embryos, we found that these criteria reliably predicted that blastoderm formation and subsequent development would proceed. We analyzed three constricted embryos that satisfied the criteria for inclusion (see Fig. 5 of the main text).

*Confocal datasets of yolk and nucleus movement:* We used these datasets to qualitatively assess the movements of yolk and nuclei. The results are shown in Fig. 1 of the main text and in Supplemental File 2.

##### 1.5 Accuracy of automated nucleus tracking

To assess the accuracy of automated nucleus tracking for lightsheet datasets, we manually tracked a set of nuclei from an embryo and compared the nucleus positions to the output from automated tracking on the same dataset. Specifically, we took the first 300 successive frames and applied the semi-automated tracking tool in MaMuT[6]. Next, we inspected each nucleus position and tracking link, and manually corrected any incorrect positions or links, resulting in 41,092 single time-point nucleus observations. Then, we used Ilastik to segment and automatically track the same dataset.

Treating the manually-corrected dataset as ground truth, we assessed the accuracy of the automated approach by calculating the distance from each nucleus in the manually-corrected dataset to the closest corresponding nucleus in the automatically-tracked dataset. Table 1 shows percentiles of these distances.

| *Percentile* | *Distance (μm)* |
|---|---|
| 10% | 0.6 |
| 25% | 0.9 |
| 50% | 1.4 |
| 75% | 1.9 |
| 90% | 2.9 |

Table 1: **Tracking discrepancy.** Percentiles of spatial discrepancies between each nucleus in a manually tracked dataset and the closest corresponding nucleus in an automatically tracked dataset. We calculated distances for 41,092 manually tracked nucleus-time points.

#### 2 Quantitative measurements of nucleus behavior

##### 2.1 Nucleus speed

For 3D+T datasets, we defined a vector

$$\vec{x}_t = (x_t, y_t, z_t) \tag{1}$$

for a nucleus position at time $t$. Using this notation, we defined instantaneous nucleus speed, $s_t$, as half of the displacement over two time points,

$$s_t = \frac{1}{2}\|\vec{x}_{t+2} - \vec{x}_t\|. \tag{2}$$

For 2D+T datasets, we defined speed in an analogous manner, calculating movement only in the $xy$ plane. The $z$-component of motion was not available in such datasets, which meant that calculated speeds are an underestimate of the true nucleus speed in 3D space.

#### 2.2 Correlation of nucleus movement vectors

For each embryo, we re-oriented the set of $\vec{x}_t$ across all $t$ so that the first principal component lay along the $x$ axis, in effect rotating the dataset so that the long axis of the embryo was parallel to the $x$ axis. By convention, we also oriented each embryo with the posterior end pointed toward the positive $x$ direction. For each $\vec{x}_t$, we computed the motion vector from $\vec{x}_t$ to $\vec{x}_{t+2}$, and then used it to calculate the correlation between pairs of nuclei moving at the same time (results shown in Fig. 1D of the main text).

#### 2.3 Local nucleus density

We defined local nucleus density around a given focal nucleus as the number of other nuclei within radius $\mathcal{R} = 150\,\mu\text{m}$.[1] For a nucleus near the periplasm of the embryo, the sphere of space within $\mathcal{R}$ included some volume that was outside of the embryo itself, but we did not wish to include this space when calculating local nucleus density. Therefore, we needed to numerically represent the surface of the eggshell so that we could exclude the volume external to the embryo from consideration in calculating local density. We took a single time point at the uniform blastoderm stage that was imaged with a multiview lightsheet microscope, segmented the nuclei, and fitted a parabola to the anterior-posterior (A-P) axis of the cloud of points. We used this parabola to transform the positions of nuclei by mapping the parabola to a straight line. Then we calculated the convex hull of the transformed points, and applied the reverse transformation to the convex hull. This produced the volume $\mathcal{B}$.

We took the sphere $\mathcal{S}$ defined by $\mathcal{R}$, and then defined

$$V = \text{volume}(\mathcal{S} \cap \mathcal{B}). \tag{3}$$

The volume fraction was computed as

$$\phi = \frac{V}{\frac{4}{3}\pi\mathcal{R}^3} \tag{4}$$

and the local density was defined as

$$\rho = \frac{\#}{\phi}, \tag{5}$$

where $\#$ represents the count of nuclei within $\mathcal{R}$.

For 2D+T datasets, local nucleus density was treated in an analogous manner: we computed the region of overlap of a 150 μm circle with the 2D convex hull of all nuclei at the blastoderm stage. Then we counted the number of nuclei within that area and calculated the area-weighted density as above. This meant that nuclear densities were

[1] We found that when we defined *local nucleus density* as the the number of other nuclei within $\mathcal{R} = 150\,\mu\text{m}$, across the range of densities found during cricket blastoderm formation, we observed a positive association between local nucleus density and mitotic cycle duration (main text Fig. 2C-E). Similarly, we observed a negative association between local nucleus density and nucleus speed (main text Figure 3E). We found that these results were not sensitive to the exact $\mathcal{R}$ we that chose. The same associations were obtained for radii from 50 to 300 μm (data not shown).

calculated from volumes with different shapes in the 3D+T vs. 2D+T datasets (i.e. spherical vs. cylindrical), and therefore their measured values cannot be compared to one another in absolute terms. Instead, we consider it appropriate to compare density-dependent results between datasets that were imaged in the same manner.

##### 2.4 Rate of change in number of nuclei

Early in cricket embryogenesis, we could not rule out the possibility that some nuclei passed through the middle of the embryo (see 1.1), where their observable fluorescence signal was sufficiently diffuse that we could not detect and track the nuclei at all. Therefore, rather than counting nuclei directly at each time point, we used *percent change in nucleus number* as a proxy for nucleus division. Because the number of detected nuclei fluctuated slightly from one time point to the next, we smoothed the total number of nuclei present by applying a Gaussian blur with a width of three time points. Using Mathematica, an interpolant, $N(t)$, was fit through the data. We differentiated this function to produce $N'(t)$. To estimate a rate of change in the number of nuclei over time, we divided $N'(t)/N(t)$. As a result, percent change was briefly calculated to be negative in the first few division cycles.

##### 2.5 Motion toward open space

We defined direction towards the "largest open space" as the vector oriented towards the centroid of the 3D Voronoi cell formed by each nucleus, bounded by the inner surface of the eggshell, as approximated by the convex hull of nuclei at the blastoderm stage.

For each nucleus, we considered $\vec{v} = \frac{1}{2}\left(\vec{x}_{t+2} - \vec{x}_t\right)$, and calculated its correlation $c$ with a vector into the direction into open space, $\vec{s}$, as

$$c = \frac{\vec{x} \cdot \vec{s}}{\|\vec{x}\| \|\vec{s}\|}. \tag{6}$$

To account for timepoint-to-timepoint noise in this calculation, we calculated average "movement into space" vectors for each nucleus over a sliding window of three time points. In addition, we computed the shortest distance from all nucleus locations $\vec{x}_t$ to the inner surface of the eggshell, $d_S$. We binned nuclei by $d_S$ into those that were near the surface ($d_S < 75\,\mu\text{m}$) and those that were far from the surface ($d_S \geq 75\,\mu\text{m}$), as shown in main text Fig. 3G,H.

#### 3 Supplemental movie of yolk and nucleus movements

We have included an example movie as Supplemental File 2. This dataset was captured as follows:

- *Genotype:* Act-H2B-EGFP; Act-mtdT
- *Subject:* The field of view was centered 20% of the way from the anterior pole of the embryo, captured during the period of syncytial development when nuclei were initially reaching the anterior pole of the embryo.

- *Z-step:* 2.5 microns
- *Duration:* 132 time steps
- *Time interval:* 2 minutes
- *$xy$ resolution:* 1 pixel = 0.327 µm
- *Image processing:* 17 consecutive z-slices were combined with as a maximal intensity projection (MIP). In order to image a given nucleus and its surrounding yolk, we only needed two to three z-slices of this thickness. We used more z-slices to capture more nuclei in a single video.

This movie illustrates that there are no obvious bulk flows of yolk, either during the phase when nuclei are moving ("Phase A") or when they are relatively immobile ("Phase B").

### 4 Simulating blastoderm formation

#### 4.1 Overview and aim

As described in the main text, the goal of our modeling approach was to assess whether a local, asymmetric, active pulling force on each nucleus could satisfactorily recapitulate the empirical patterns of nucleus behaviors. Here we report on the model in closer detail. The simulation is performed in a $202 \times 92 \times 92$ volume region. We verified results are consistent with scaling up the grid by a factor of two, but for time efficiency we opted to use the smaller grid. The simulation time step is $T = 45$ s corresponding to half a time step from the lightsheet data. The embryo used for the fit of the simulation boundary was 2421 µm long, and thus the grid spacing is $L = 11.98$ µm.

#### 4.2 Model components

We simulated the center of each nucleus $\vec{x}_i(t)$ as a function of time. The nucleus positions followed the differential equation

$$\alpha M \frac{d^2\vec{x}_i}{dt^2} + \gamma \left( \frac{d\vec{x}_i}{dt} - \vec{v}^{cyt}(\vec{x}_i(t)) \right) = \vec{F}_i^{tot} \tag{7}$$

where $M$ is an arbitrary mass scale and $\alpha M$ is the mass of a nucleus, $\gamma$ is a drag coefficient between the nucleus and the cytoplasm, and $\vec{F}_i^{tot}$ is the sum total of forces on the nucleus. Based on our experimental results, we assumed that $\vec{v}^{cyt} = \vec{0}$ and we did not simulate any viscous or viscoelastic flow in the cytoplasm. We worked in the overdamped limit where the acceleration could be neglected, yielding

$$\frac{d\vec{x}_i}{dt} = \frac{1}{\gamma} \vec{F}_i^{tot}, \tag{8}$$

We simulated equation (8) using the first-order forward Euler method. We describe the different contributions to the force below. We take $\gamma = 1M/T = 0.022M$ s$^{-1}$.

###### 4.2.1 Embryo

We used the shape of a real cricket embryo, determined from the positions of nuclei during the uniform blastoderm stage, as described in Section 2.3. We approximated each cross section along the A-P axis $z$ with an ellipse with radii $r_1(z)$ and $r_2(z)$ with a center at $(x(z), y(z))$. This defined a boundary $\mathcal{B}$. For simulated *G. bimaculatus* embryos, we performed all calculations on the discretized 3D grid. The grid was used by the scikit-fmm library, which we used to compute the clouds for each nucleus.

###### 4.2.2 Division geometry

Nuclei divided in random directions (i.e. with random spindle orientation) throughout the simulation, irrespective of the proximity of neighbors, the proximity of the eggshell, or the orientation of previous divisions in a lineage. Newly divided daughter nuclei were created at $\vec{x}^{mother} \pm \vec{n}(1\,\mu\text{m})$, where $\vec{n}$ is a random unit normal. If this displacement resulted in a daughter nucleus landing outside the boundary of the embryo, we assigned a new position for the nucleus by projecting it back to the surface of the embryo boundary by finding the closest point on the surface.

###### 4.2.3 Density-dependent cell cycle duration

In the empirical data, we observed a strong positive association between local nucleus density and cell cycle duration (see Fig. 2 in the main text). We incorporated this relationship into our simulation directly. We fitted a logistic curve to the empirical density vs. cell cycle duration data (shown in Fig. 2 in the main text), and then drew from it with normally distributed noise, based on the density at nucleus birth. In the real embryos, we observed that the variation in cycle duration increased as density increased (Fig. 2C-E), so we fitted the variance to a global clock used in the simulation, $(90\text{ s}) + (270\text{ s})(t - T)/T$, where $T$ was the total simulation time.

###### 4.2.4 Asymmetric pulling cloud

We hypothesized that there was a local, asymmetric, active pulling force on each nucleus. We describe the biological inspiration for this model in the main text. Here we describe the implementation of the model.

Our force calculation models the nuclei as spheres with a 7.2 µm diameter. We modeled the motion of a nucleus by ascribing a pulling cloud that emerges from a single origin on the surface of the nucleus. The cloud grows uniformly in all directions to a maximal radius, $R_{\text{max}} = 150\,\mu\text{m}$, except for where its growth is impeded by the surface of the nucleus from which it originated, the surface of another cloud region, or the inner surface of the eggshell.

For each nucleus, we computed this cloud region by using the Python library scikit-fmm[10]. We used a fast marching method[11], which is a computationally efficient way to identify all the voxels in the discretized grid that should be assigned to each nucleus. Specifically, we initialized the locations of nuclei to be zeroes of a field $T(\vec{x})$. We then solved the Eikonal equation

$$|\nabla T(\vec{x})| = 1, \tag{9}$$

where $T(\vec{x})$ describes the minimum travel time from a nucleus to each position. The fast marching method works by scanning outward from the nuclei positions to fill in the values of the $T(\vec{x})$ on the discretized grid. It also keeps track of which nucleus is closest to each point $\vec{x}$. This allows us to efficiently compute the region $\mathcal{R}_i$ that is closest to each nucleus $\vec{x}_i$. From here, we define the pulling cloud region $\mathcal{V}_i$ by removing space that is further away than $R_{\text{max}}$, and space that is occluded by the steric effects from the nucleus (*i.e.* where a ray from the cloud origin to $\vec{x}$ intersects the nucleus).

Each point in the cloud region creates a tug toward its position. Since our model is based on the assumption that this tug arises from a set of microtubules emanating from the cloud origin, we scale the tug by a factor of $1/r^2$ to account for the microtubule density, where $r$ is the distance to the nucleus center.[2] Hence, the total force created by the cloud is given by the integral

$$\vec{F}_i^{cloud} = \int_{\mathcal{V}_i} \beta \frac{\vec{y} - \vec{x}_i}{\|\vec{y} - \vec{x}_i\|_2^3} d\vec{y}. \tag{10}$$

The parameter $\beta = 0.00521 M/T^2 = 2.57 \times 10^{-6}\ M\,\text{s}^{-2}$ was fit from experimental data where we matched the maximum speed at the 4-nucleus stage. The integral in Eq. (10) is performed by summing over all voxels in $\mathcal{V}_i$.

###### 4.2.5 Bias of nucleus movement toward periplasm

In *G. bimaculatus* embryos, the first zygotic division tends to occur about ~60% of the way along the A-P axis, as measured from the anterior pole of the embryo. During the first few cell cycles, nuclei spread apart in space, with some of them moving through "open space" across the yolk-rich middle of the embryo, and others moving within the periplasm, close to the inner surface of the eggshell. The nuclei ultimately move into the periplasm and stay there (see Fig. 1A, B). Because nuclei travel varied paths and arrive in the periplasm asynchronously, we model their movement into the periplasm as a small bias that contributes additively to their movement. Specifically, for each nucleus $\vec{x}_i$ we introduced a component to a nucleus's motion vector that is in the direction of the nearest point $\vec{y}_i$ on the inner surface of the eggshell. We define the vector $\vec{q}_i = \vec{y}_i - \vec{x}_i$. The contribution to the force is given by

$$\vec{F}_i^{surface} = \zeta \|\vec{F}_i^{cloud}\| \frac{\vec{q}_i}{\|\vec{q}_i\|} \tag{11}$$

where $\zeta$ is a dimensionless constant. We include a factor $\|\vec{F}_i^{cloud}\|$ since it is an existing force scale in the model. We used a value of $\zeta = \frac{1}{3}$ to match the bias towards the periplasm in the empirical data.

[2] Technically it would be more consistent with the model to normalize by $1/\tilde{r}^2$, where $\tilde{r}$ is the distance to the cloud origin on the nucleus surface. However, this could create numerical difficulties since the cloud origin is on the boundary of $\mathcal{V}_i$ and thus $\tilde{r}$ can become arbitrarily small. Since the cloud origin and nucleus center are close, using $r$ instead of $\tilde{r}$ has a minimal effect on the outcome and is more numerically stable.

##### 4.3 Assessing a model of mutual repulsion

We replaced our calculation of cloud-based pulling forces with the mutual repulsion forces presented by Dutta and colleagues[12], which was tuned on a later stage of *Drosophila melanogaster* development. With this model, there is a decay in nuclear speed as nuclei separate from one another (see Equation 2[12], data not shown). This is the opposite of what we observe in real *G. bimaculatus* embryos and in simulations that use pulling clouds. In both cases, nuclei that move fastest are those that are farthest from other nuclei. Therefore, we concluded that a mechanism based on mutual repulsion does not effectively explain nucleus movements in *G. bimaculatus*.

#### 5 Methods for physically constricting embryos

##### 5.1 Assembling the embryo constriction device

We constricted embryos in a custom device, constructed as follows: We used a laser cutter to cut the components from sheets of acrylic (thickness = 1.59 mm, McMaster-Carr 8560K172; thickness = 3.18 mm, McMaster-Carr 8560K257), and then assembled them with acrylic welding solution (IPS Weld-On 3 Acrylic Plastic Cement) following the procedure described previously[7]. A DXF design file of the acrylic components is included as Supplementary File 3. The device's construction also required a common binder clip (width = 20 mm), two steel nails (diameter = 2 mm; length = 39 mm), and a rubber band (thickness = 1 mm). We were able to successfully constrict embryos with versions of the device that were assembled with several different types of nails, rubber bands, and binder clips (data not shown). We used a human hair as the constricting fiber. It was held in place by being pinched in a block of elastomer (Dow Corning Sylgard 184 Elastomer Kit) into which a slit had been cut with a razor blade. We used a crafting hot glue gun to attach acrylic components to non-acrylic components. Schematics of all components are shown to scale along with additional assembly instructions in Fig. S1.

##### 5.2 Using the embryo constriction device

Figure S2A is a photograph of the device in use, with human hairs used as the constricting fibers. We placed embryos in water-filled acrylic troughs on a removable platform, and then constricted them one at a time. To do so, we threaded a hair through the removable plastic platform, around an embryo, back through a hole at the bottom of a trough in the removable platform, and then attached it to a ratchet mechanism. Detailed instructions for this procedure are shown in Fig. S2B. This allowed us to increase tension on the hair while observing the embryo under a dissection microscope. As we increased tension, the embryo was incrementally constricted. After the desired extent of constriction was achieved, we placed temporary dabs of hot glue to affix the hair in place for the duration of imaging.

The device was able to hold multiple constricted embryos at a time for simultaneous imaging, up to a maximum of seven embryos. Once a set of embryos was constricted, we took the removable platform from the constriction device, placed it in a glass bottom 6-well dish (MatTek P06G-1.5-20-F), and imaged it using a Zeiss Cell Discoverer microscope following previously described methods[7].

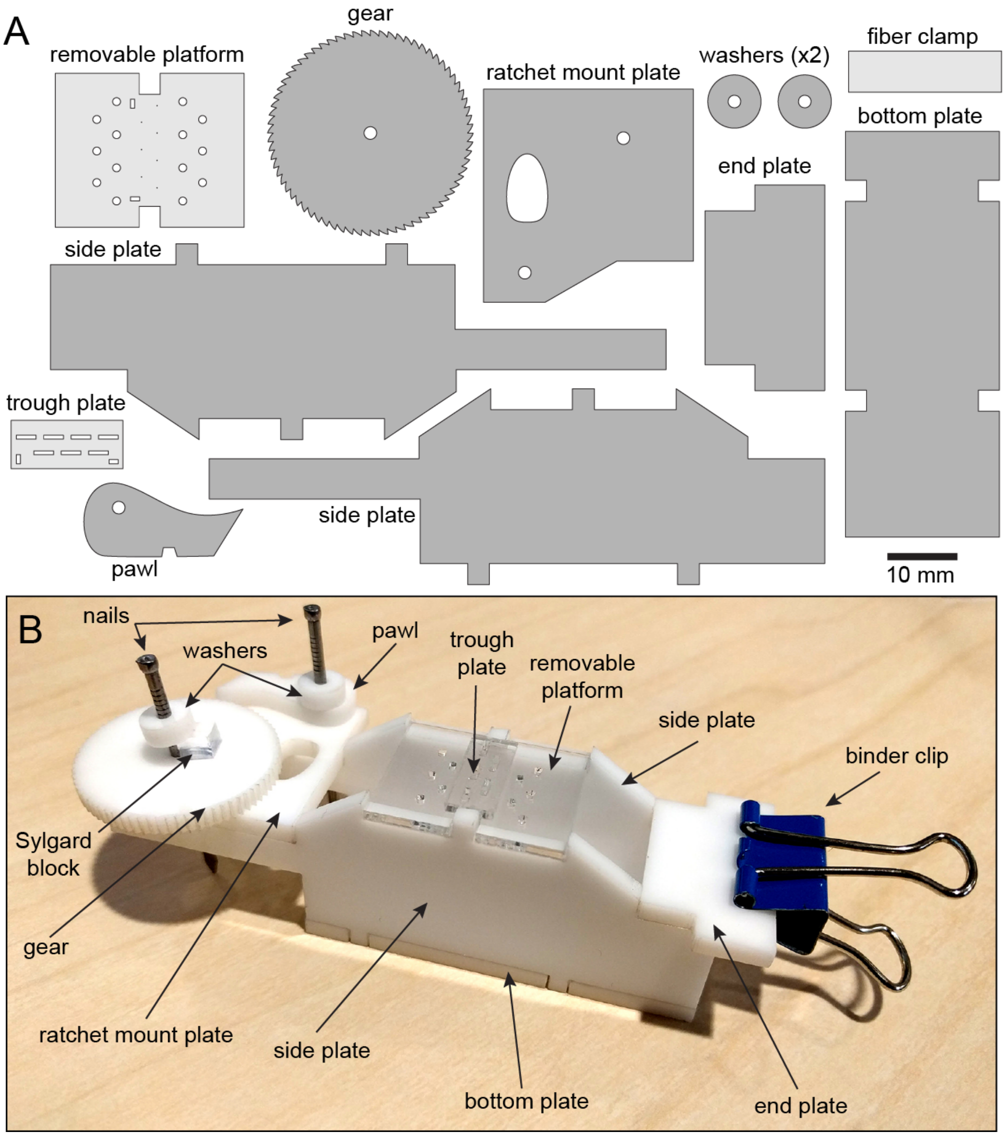

Figure S1: **Assembling a device for constricting embryos**. **A**, Diagram of all acrylic components. We cut the light gray pieces from 1.59 mm-thick acrylic sheet. We cut the rest from 3.18 mm-thick acrylic sheet. **B**, The assembled device also included a binder clip and two steel nails. Before welding the trough plate to the removable platform, we positioned trough plate so that the rectangular holes in the platform lined up with the corresponding holes in the trough plate. After the rest of the components were welded together as shown, we inserted one steel nail through a washer, then through the pawl, then through the ratchet mount plate. We applied hot glue to the top of the washer, fixing it to the nail. We also applied hot glue to the underside of the ratchet mount plate where the nail emerged, fixing the nail to the plate. We used a straight edge razor to cut a block of Sylgard elastomer with approximate dimensions 4 mm $\times$ 8 mm $\times$ 8 mm, and then sliced the block through half its depth when oriented with the largest face flat on a table. We inserted the second nail through the other washer, gear, and then the ratchet mount plate. We positioned the Sylgard block as shown, and then hot glued to the gear. We applied hot glue to the top of the second washer, fixing it to the nail. Finally, we applied more hot glue to the nail where it emerged underneath the ratchet mount plate, fixing it in place.

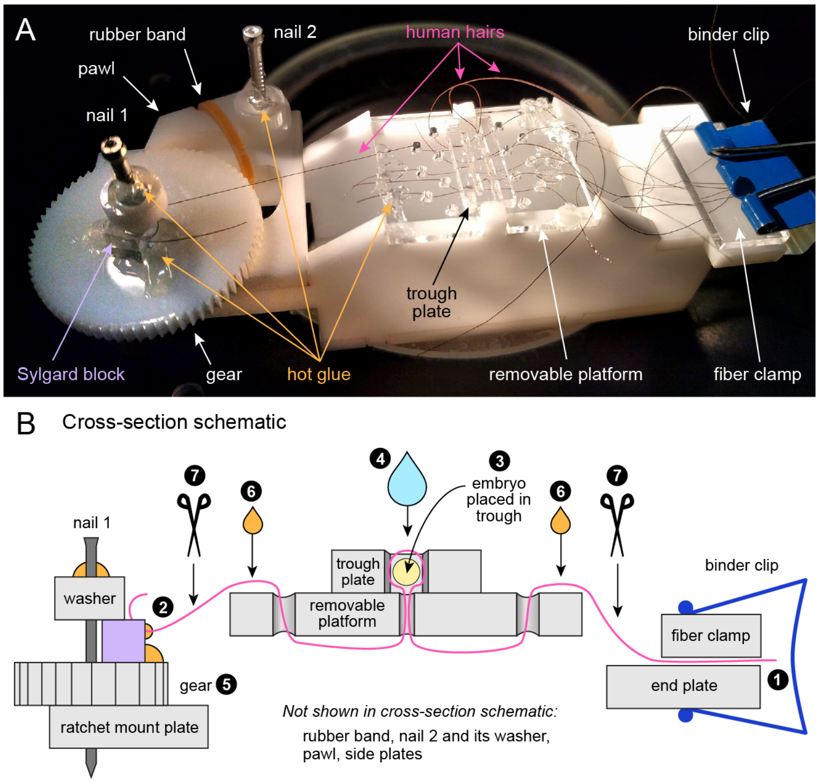

Figure S2: **Constricting embryos.** **A**, Photograph of the constriction device in use on a dissection microscope. To prepare the device for use, we cut a small rubber band to linearize it, inserted it through the largest hole in the ratchet mount plate, wrapped it underneath the side plate, and then rejoined it to itself by tying a knot. The re-joined rubber band formed a loop with the band slotting into the notch on the pawl, as shown. The tension on the rubber band pushed the pawl against the gear to make a ratchet. White arrows highlight the rubber band, binder clip, fiber clamp, removable platform, and gear. Hot glue is indicated with orange arrows. Note: the hot glue, rubber band, and fiber clamp were not shown in Fig. S1. The fiber clamp was an unattached rectangle of acrylic that, when squeezed by the binder clip, effectively held the fibers clamped in place. **B**, Cut-away schematic of fiber threading path (not to scale). Acrylic components are depicted in light gray, nail in dark gray, Sylgard block in lavender, hot glue in orange, water in light blue, and binder clip in dark blue. The embryo is shown in yellow as a cross-section end-on. The constricting fiber (a human hair) is shown in pink.

1. We clamped one hair between the end plate and fiber clamp.
2. We threaded the hair by hand through the path shown, and ultimately inserted into it the slit in the elastomer block, leaving a large loop in the place around the trough where the embryo would go. The removable platform can be taken off of the device to make threading easier.
3. We placed the embryo into the trough.
4. We added distilled water to the trough to fill its entire volume.
5. We manually cranked the gear while the embryo was observed by the user with a dissection microscope. Surface tension kept the water from leaking through the bottom hole.
6. When the desired constriction was achieved, we used hot glue to affix the hair at the two locations indicated with orange droplets.
7. We used scissors to cut the hair at the two locations indicated with the scissors icons. We repeated the constriction process on multiple embryos and then removed the removable plate. To image the embryos, we inverted the removable plate, with constricted embryos still in the troughs, and placed it into a coverslip-bottomed dish. We filled the dish with distilled water, and then imaged the embryos with an inverted microscope.

### 6 Code, data, and materials availability

A GitHub repository (https://github.com/hoffmannjordan/gryllus_nuclear_movements) has the code used to simulate nuclear movements during blastoderm formation and the code used to convert the tracking output from Ilastik into an XML file that was parsable by MaMuT. Plots were generated with Mathematica; notebooks used to generate each type of data visualization are available from the corresponding authors upon request.

A dataset of 3D+T nucleus positions and tracking links that supports the findings of this study is available at the GitHub repository for this project as well. Additional data that support the findings of this study are available from the corresponding authors upon reasonable request.

The *G. bimaculatus* culture is available for sharing from the corresponding authors upon request, provided that the requestor obtains the necessary permits for the transfer and continued maintenance of the culture (the specific permits will vary by jurisdiction). The plasmid for generating the Act-mtdT transgenic line described in the present study will likewise be available from the corresponding authors upon reasonable request.
